## Supplementary material for "Conserved nuclear receptors controlling a novel trait target fast-evolving genes expressed in a single cell": All supplemental figures and tables

**Supplementary figures**

**SFig. 1.** Additional images. (**A**) The mouth of the wild-type eurystomatous (Eu) morph, wild-type stenostomatous (St) morph, *nhr-1* mutant, and *nhr-40* mutant in two focal planes. (**B**) Expression pattern of an *nhr-1* transcriptional reporter in a young larva. TurboRFP channel is presented as a maximum intensity projection. (**C**) Antibody staining against the HA epitope in a line, in which the tag was “knocked in” into the endogenous locus. Fluorescent channel is presented as a maximum intensity projection. (**D**) Expression patterns of *nhr-40* and *nhr-1* transcriptional reporters in a double reporter line. TurboRFP (magenta) and Venus (green) channels are presented as standard deviation and maximum intensity projections, respectively. Co-expression results in white color. (**E**) Expression patterns of *nhr-40* and *nhr-1* transcriptional reporters in a double reporter line. TurboRFP is encoded as magenta, Venus as green. Co-expression results in white color. D = dorsal, V= ventral, A = anterior, P = posterior.

**SFig. 2.** Additional bioinformatic analyses. (**A**) Whole-genome re-sequencing and RNA-seq of the *null* allele of *nhr-40*. (**B**) Bulk segregant analysis of the suppressor of *nhr-40(tu505)* with the location of *nhr-1* marked with a dotted line. See STable 1 for the list of non-synonymous and nonsense substitutions within the candidate region.

**SFig. 3.** Geometric morphometric analysis of 20 landmarks in the mouth of the wild-type strain RS2333, the *nhr-1(tu1163)* mutant isolated in this study, and the *daf-21*/Hsp90*(tu519)* mutant previously shown to exhibit an aberrant mouth morphology while maintaining the dimorphism. Each point in the PCA plot corresponds to a single animal. Deformation grids at the extremes of the two axes display differences to the mean shape of all individuals.

**STable 1.** List of non-synonymous and nonsense substitutions within the candidate region on chromosome X identified through the bulk segregant analysis of the suppressor of *nhr-40(tu505)*.

**STable 2.** Description of transgenic constructs. Promoters include 5' UTRs and may include coding exons, and introns. See SData 1 for the sequences of the listed elements. CDS = coding sequence, UTR = untranslated sequence.

**SData 1.** FASTA file of nucleotide sequences used to create transgenic constructs.

**SData 2.** Phylogenetic trees from Fig. 5 and alignments used to generate them.

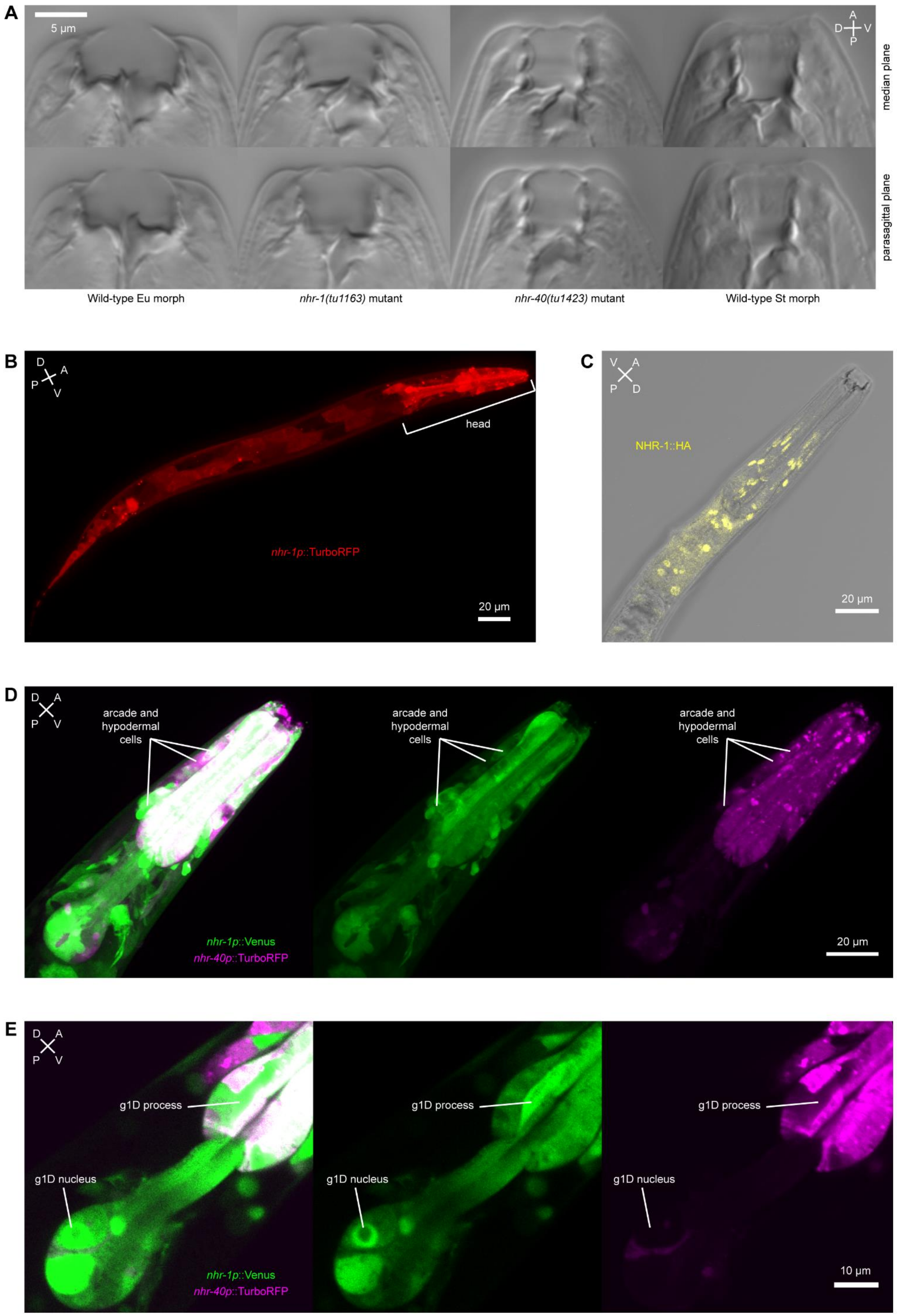
A
5 µm
A
D
V
P
median plane
parasagittal plane
Wild-type Eu morph
nhr-1(tu1163) mutant
nhr-40(tu1423) mutant
Wild-type St morph
B
D
A
P
V
head
nhr-1p::TurboRFP
20 µm
C
V
A
P
D
NHR-1::HA
20 µm
D
D
A
P
V
arcade and hypodermal cells
arcade and hypodermal cells
arcade and hypodermal cells
nhr-1p::Venus
nhr-40p::TurboRFP
20 µm
E
D
A
P
V
g1D process
g1D process
g1D process
g1D nucleus
g1D nucleus
g1D nucleus
nhr-1p::Venus
nhr-40p::TurboRFP
10 µm

**SFig. 1.**

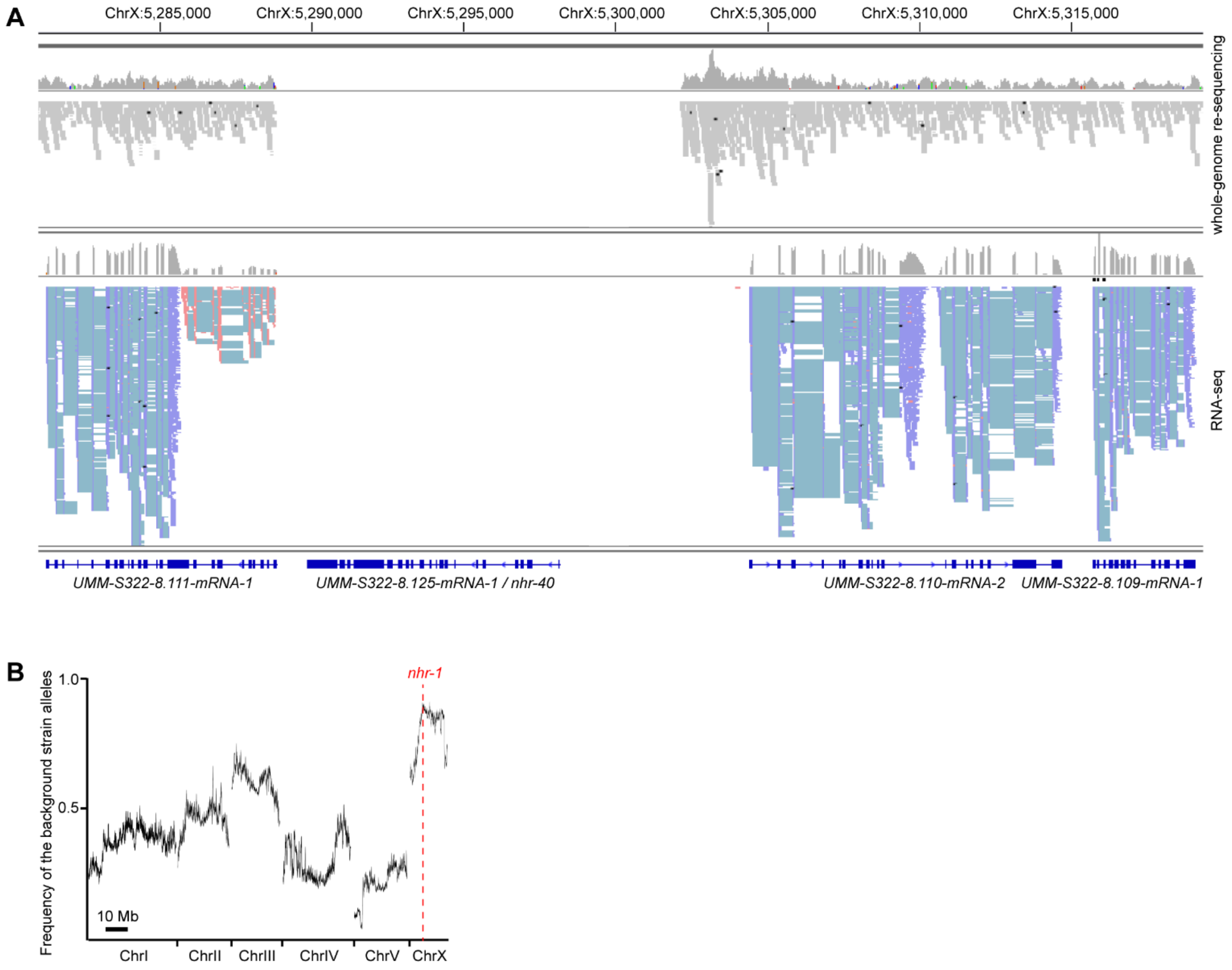
A
ChrX:5,285,000
ChrX:5,290,000
ChrX:5,295,000
ChrX:5,300,000
ChrX:5,305,000
ChrX:5,310,000
ChrX:5,315,000
whole-genome re-sequencing
RNA-seq
UMM-S322-8.111-mRNA-1
UMM-S322-8.125-mRNA-1 / nhr-40
UMM-S322-8.110-mRNA-2
UMM-S322-8.109-mRNA-1
B
nhr-1
1.0
0.5
Frequency of the background strain alleles
10 Mb
ChrI
ChrII
ChrIII
ChrIV
ChrV
ChrX

**SFig. 2.**

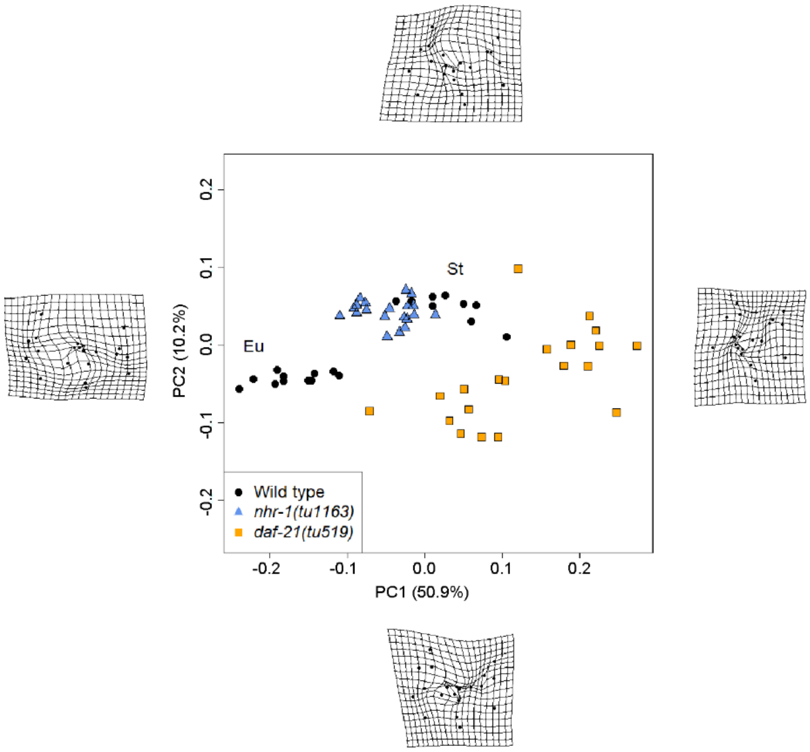

St
Eu
PC2 (10.2%)
PC1 (50.9%)
0.2
0.1
0.0
-0.1
-0.2
-0.2
-0.1
0.0
0.1
0.2
Wild type
nhr-1(tu1163)
daf-21(tu519)


**SFig. 3.**

STable 1

| Chromosome | Position | Ref allele | Mutant allele | Wormbase WS268 Identifier | El Paco annotation v1 Identifier | Putative *C. elegans* ortholog | Classification |
|---|---|---|---|---|---|---|---|
| ChrX | 5699852 | A | T | *PPA39635* | *UMM-S322-4.13-mRNA-1* | | non-synonymous |
| ChrX | 5753767 | C | T | *PPA21915* | *UMM-S322-4.28-mRNA-1* | *nhr-1* | non-synonymous |
| ChrX | 6299273 | C | T | *PPA10517* | *UMM-S429-1.43-mRNA-1* | *glr-8* | non-synonymous |
| ChrX | 6367712 | A | C | *PPA40141* | *UMM-S429-2.95-mRNA-1* | | non-synonymous |
| ChrX | 6380013 | A | T | *PPA05325* | *UMM-S429-2.97-mRNA-1* | *nep-17* | non-synonymous |
| ChrX | 6402673 | C | T | *PPA40146* | *UMM-S429-2.115-mRNA-1* | | non-synonymous |
| ChrX | 6443259 | T | A | *PPA05313* | *UMM-S429-2.24-mRNA-1* | *K09E2.1* | non-synonymous |
| ChrX | 6444545 | G | C | *PPA05313* | *UMM-S429-2.24-mRNA-1* | *K09E2.1* | non-synonymous |
| ChrX | 6957752 | C | T | *PPA18712* | *UMM-S293-2.13-mRNA-1* | *sec-3* | non-synonymous |
| ChrX | 7145062 | C | T | *PPA39440* | *UMM-S293-4.13-mRNA-1* | | non-synonymous |
| ChrX | 7479342 | T | A | *PPA18845* | *UMM-S293-7.70-mRNA-1* | *ZK470.2* | non-synonymous |
| ChrX | 7479349 | C | A | *PPA18845* | *UMM-S293-7.70-mRNA-1* | *ZK470.2* | non-synonymous |
| ChrX | 7900717 | A | G | *PPA09604* | *UMM-S293-11.35-mRNA-1* | | non-synonymous |
| ChrX | 8602408 | C | T | *PPA45298* | *UMS-S328-6.31-mRNA-1* | | non-synonymous |
| ChrX | 8866724 | C | T | *PPA39709* | *UMM-S328-3.19-mRNA-1* | | non-synonymous |
| ChrX | 9098848 | T | G | *PPA30108* | *UMS-S328-0.4-mRNA-1* | | non-synonymous |
| ChrX | 9098849 | T | A | *PPA30108* | *UMS-S328-0.4-mRNA-1* | | nonsense |
| ChrX | 9099400 | A | G | *PPA30108* | *UMS-S328-0.4-mRNA-1* | | non-synonymous |
| ChrX | 9387377 | C | T | *PPA07538* | *UMM-S419-3.4-mRNA-1* | *flp-11* | non-synonymous |
| ChrX | 9894810 | G | A | *PPA20385* | *UMM-S2857-8.19-mRNA-1* | *tap-1* | non-synonymous |
| ChrX | 1.2E+07 | C | T | *PPA03807* | *UMM-S2859-11.66-mRNA-3* | | non-synonymous |
| ChrX | 1.4E+07 | A | T | *PPA39997* | *UMM-S398-3.15-mRNA-1* | | non-synonymous |
| ChrX | 1.4E+07 | G | A | *PPA40003* | *UMM-S398-3.10-mRNA-1* | | non-synonymous |
| ChrX | 1.4E+07 | G | A | *PPA40012* | *UMM-S398-4.109-mRNA-1* | | non-synonymous |
| ChrX | 1.4E+07 | A | C | *PPA40020* | *UMM-S398-4.137-mRNA-1* | *gcy-36* | non-synonymous |
| ChrX | 1.4E+07 | A | G | *PPA42287* | *UMM-S2845-2.23-mRNA-1* | | non-synonymous |
| ChrX | 1.4E+07 | A | T | *PPA42287* | *UMM-S2845-2.23-mRNA-1* | | non-synonymous |
| ChrX | 1.6E+07 | A | G | *PPA42394* | *UMM-S2845-19.65-mRNA-1* | | non-synonymous |

STable 2

| Construct | Plasmid | Promoter | CDS | 3' UTR |
|---|---|---|---|---|
| *nhr-1* rescue construct | pBS14 | *nhr-1* (short) | *nhr-1* , with synthetic introns | *nhr-1* |
| HA-tagged *nhr-1* rescue construct | pBS25 | *nhr-1* (short) | *nhr-1* , with synthetic introns and C-terminal HA tag | *nhr-1* |
| red *nhr-1* reporter | pBS9 | *nhr-1* (long) | TurboRFP | *rpl-23* |
| yellow *nhr-1* reporter | pBS36 | *nhr-1* (long) | Venus | *rpl-23* |
| red *nhr-40* reporter | pBS30 | *nhr-40* | TurboRFP | *rpl-23* |
| red *UMA-S293-8.46-mRNA-1* reporter | pSS1 | *UMA-S293-8.46-mRNA-1* | TurboRFP | *rpl-23* |
| red *UMM-S2847-6.45-mRNA-1* reporter | pSS2 | *UMM-S2847-6.45-mRNA-1* | TurboRFP | *rpl-23* |
| red *UMM-S2857-0.30-mRNA-1* reporter | pSS6 | *UMM-S2857-0.30-mRNA-1* | TurboRFP | *rpl-23* |
| red *UMS-S2861-1.50-mRNA-1* reporter | pSS4 | *UMS-S2861-1.50-mRNA-1* | TurboRFP | *rpl-23* |
| red *UMM-S328-9.28-mRNA-1* reporter | pSS3 | *UMM-S328-9.28-mRNA-1* | TurboRFP | *rpl-23* |
| red *UMM-S2857-0.41-mRNA-1* reporter | pSS8 | *UMM-S2857-0.41-mRNA-1* | TurboRFP | *rpl-23* |
| red *UMM-S283-11.38-mRNA-1* reporter | pBS26 | *UMM-S283-11.38-mRNA-1* | TurboRFP | *rpl-23* |
| red *UMM-S328-10.33-mRNA-1* reporter | pBS27 | *UMM-S328-10.33-mRNA-1* | TurboRFP | *rpl-23* |
| red *UMS-S328-0.4-mRNA-1* reporter | pBS28 | *UMS-S328-0.4-mRNA-1* | TurboRFP | *rpl-23* |
